# Meta-Analysis Reveals That Explore-Exploit Decisions are Dissociable by Activation in the Dorsal Lateral Prefrontal Cortex, Anterior Insula, and the Dorsal Anterior Cingulate Cortex

**DOI:** 10.1101/2023.10.21.563317

**Authors:** Daniel Sazhin, Abraham Dachs, David V. Smith

## Abstract

Explore-exploit research faces challenges in generalizability due to a limited theoretical basis for exploration and exploitation. Neuroimaging can help identify whether explore-exploit decisions involve an opponent processing system to address this issue. Thus, we conducted a coordinate-based meta-analysis (N=23 studies) finding activation in the dorsal lateral prefrontal cortex, anterior insula, and anterior cingulate cortex during exploration versus exploitation, which provides some evidence for opponent processing. However, the conjunction of explore-exploit decisions was associated with activation in the dorsal anterior cingulate cortex and dorsal medial prefrontal cortex, suggesting that these brain regions do not engage in opponent processing. Furthermore, exploratory analyses revealed heterogeneity in brain responses between task types during exploration and exploitation respectively. Coupled with results suggesting that activation during exploration and exploitation decisions is generally more similar than it is different suggests that there remain significant challenges in characterizing explore-exploit decision making. Nonetheless, dorsal lateral prefrontal cortex, anterior insula, and dorsal anterior cingulate cortex activation differentiate explore and exploit decisions and identifying these responses can aid in targeted interventions aimed at manipulating these decisions.

Graphical Abstract
We conducted a coordinate-based meta-analysis (N=23 studies) where we found activation in the dorsal lateral prefrontal cortex, anterior insula, and the anterior cingulate cortex during exploration versus exploitation. However, the conjunction of explore-exploit decisions was associated with activation in the dorsal medial prefrontal cortex, and the insula, suggesting that these brain regions do not engage in opponent processing. Nonetheless, activation that differentiates explore and exploit decisions and can help in targeted interventions aimed at manipulating these decisions.

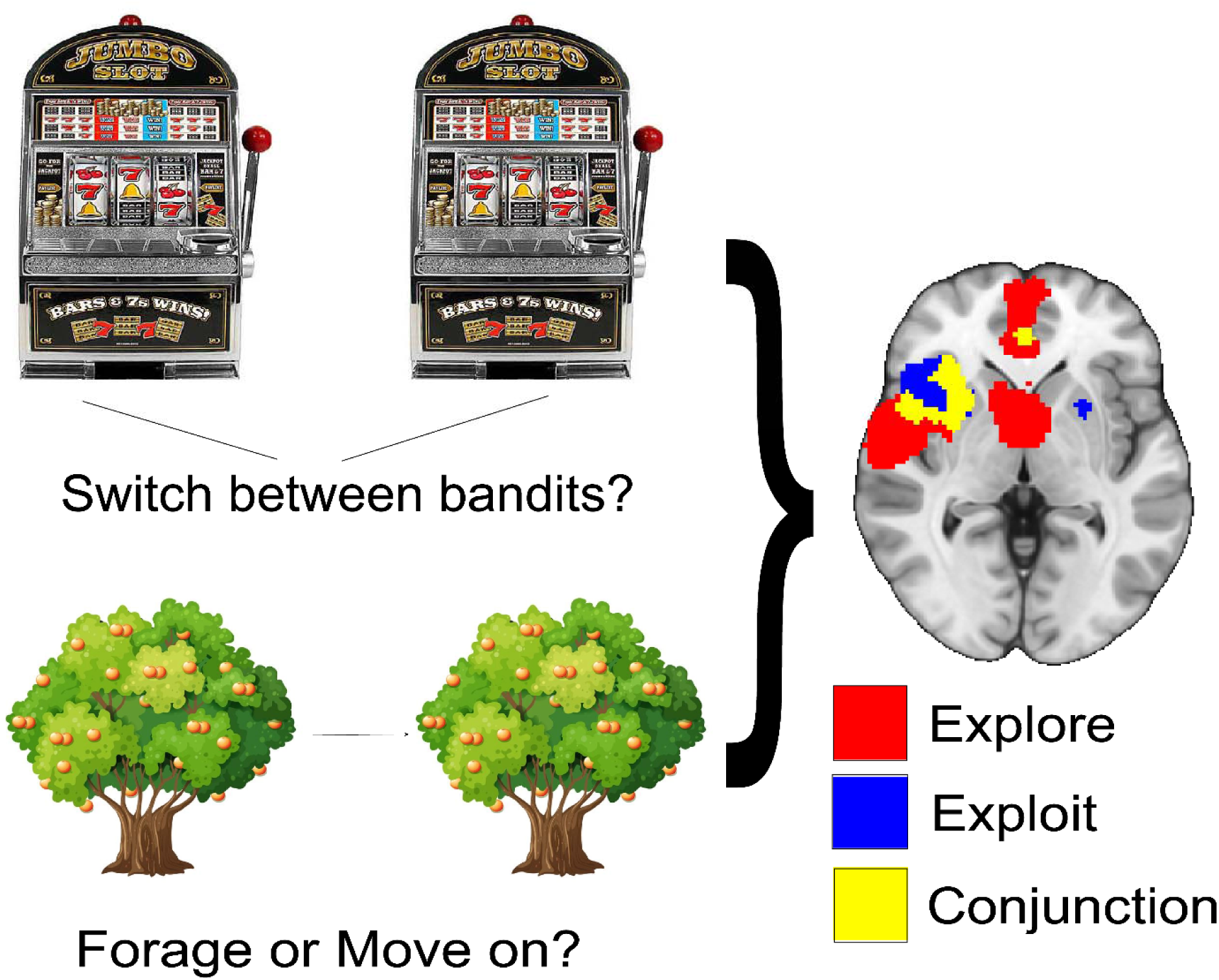

## 1. Introduction

Explore-exploit problems are ubiquitous in many real-world situations such as staying in one line of employment or moving to another, keeping versus selling a stock, or trying out a new ice cream flavor versus sticking with what you know. In situations where a person does not have full knowledge of their opportunities and outcomes, there is a fundamental dilemma of whether to explore the space of possibilities available to them or to exploit what they already know. Due to their prevalence in naturalistic settings, explore-exploit dilemmas have been extensively investigated, with an emphasis on whether certain people are consistently likely to overexploit or underexploit. Over and under exploitation is especially interesting in psychological research as markers of psychopathology, such as among people with anxiety, compulsivity and smoking habits (Aberg and Paz 2022; Merideth A. Addicott et al. 2014; L. S. Morris et al. 2016).

Despite the interest in explore-exploit tasks, generating generalizable insights from decisions made in the lab presents several major challenges. The first challenge is that explore-exploit situations generally involve many independent variables that are difficult to control, such as the hidden payoffs of existing options, the number of options available to the participant, the strategies guiding exploration (random or directed), and the time horizons of the tasks (Wilson et al. 2014). Even simply understanding the payoffs of these choices include a multitude of decision variables such as risk, uncertainty, and ambiguity. Overall, the independent variables investigated, such as uncertainty (Payzan-LeNestour et al. 2013), task difficulty (Lee and Daunizeau 2021), information search (Drugowitsch et al. 2012) suggest explore-exploit decisions are a subset of within a value-based decision-making process. Controlling these independent variables is necessary to assess if exploration and exploitation can be construed as a consistent and useful psychological construct.

Second, is a lack of behavioral convergence across foraging and n-armed bandit tasks (von Helversen et al. 2018), which suggests that exploration and exploitation may not be guided by consistent attitudes. Third, there remains a lack of a unified theory of exploration and exploitation, behaviorally and neurally, as to whether exploration and exploitation are opponent processes, or the result of the interaction of multiple underlying systems. These major questions suggest that reviewing the common features of exploration and exploitation could yield clarity both theoretically and empirically regarding how the field should understand these decisions. Specifically, by assessing common and distinct patterns of activation between exploration and exploitation across n-armed bandit and foraging tasks, it may be possible to identify evidence for whether explore-exploit decisions are dissociable psychological constructs. To do so, we first review extant literature, followed by conducting a coordinate-based (CBMA) meta-analysis to understand which brain regions are involved in exploration and exploitation.

### 1.1 Understanding Explore-Exploit Decisions Behaviorally

Many tasks have been conceived to isolate explore-exploit decisions, though they mostly fall within two categories: foraging tasks (M. A. Addicott et al. 2017) and n-armed bandit tasks (Cohen, McClure, and Yu 2007; M. A. Addicott et al. 2017; Zhen et al. 2022). These tasks are highly prevalent in explore-exploit research because they have computationally optimal closed-form solutions. In foraging tasks, a participant selects whether to forage from a patch of resources such as an apple tree, or to travel to another patch at some distance from the current patch (Mobbs et al. 2018; Bernstein, Kacelnik, and Krebs 1988; Stephens and Krebs 1986). The optimal strategy is determined by a marginal value theorem (pMVT) which is based on the payoffs within a current patch and the distance to the next patch (Charnov 1976). Foraging tasks can be modeled through Markov decision processes (Averbeck 2015). In n-armed bandit tasks, the participant decides which slot machine they would like to sample from (Bellman and Kalaba 1957). Explore-exploit decisions are classified through a variety of computational algorithms, such as Boltzmann exploration (softmax), reinforcement learning (Kuleshov and Precup 2000), and can be approximated through Partially Observable Markov Decision Processes (POMDP) (Krishnamurthy and Wahlberg 2009). Ultimately, when the participant chooses bandits higher expected value, the decisions are classified as exploitative and when they choose bandits with lower or unknown expected value, they are classified as explorative (Daw et al. 2006).

While there are canonical foraging and n-armed bandit tasks, there are many other variants of these tasks. One variation of the n-armed bandit task is the Horizon Task which runs for 15, 30, or 45 minutes and was developed to discern if task length affects behavior (Trudel et al. 2020). Another variation of the n-armed bandit is the Leapfrog task where two bandits’ values are fixed until the lower value bandit ‘leapfrogs’ over the higher value bandit at random intervals (Blanco et al. 2016). Variations of foraging also include the Clock Task (Moustafa et al. 2008). Optimal stopping problems such as the Secretary Task (Freeman 1983) are also sometimes grouped as explore-exploit dilemmas. With such a variety of tasks, a critical question is whether the independent variables manipulated within these tasks guide exploration and exploitation, or if general tastes in exploration and exploitation tend to guide behavior. If choices are inconsistent between tasks, then exploration and exploitation should not be conceived as independent constructs, but rather as the interaction of the underlying independent variables.

Recent evidence suggests that foraging tasks and n-armed bandits lack behavioral convergence (von Helversen et al. 2018) which suggests that how people explore and exploit in n-armed bandits does not predict how people will explore or exploit in a foraging task. The lack of behavioral convergence between tasks is a major challenge as this suggests that there is a lack of a unifying psychological mechanism underlying exploration and exploitation decisions.

Another approach may be to assess if economic or psychological differences can reliably differences in exploration or exploitation. Investigators have found that the explore-exploit tradeoff was associated information gain and the level of recent rewards (Cogliati Dezza et al. 2017), and that this effect was modulated based on the cognitive load experienced by the participant (Cogliati Dezza, Cleeremans, and Alexander 2019). In the context of temporal discounting problems there have been mixed findings, with one investigation finding associations between temporal discounting and directed exploration and no relationship between temporal discounting and random exploration (Sadeghiyeh et al. 2020) and another study suggesting inconsistent preferences for temporal discounting and exploration and exploitation across multiple studies (Meyers and Koehler 2021). In assessing effects of impulsive behaviors, or risk attitudes, there were no significant associations with foraging decisions though gamblers exhibited more exploratory behavior (Merideth A. Addicott et al. 2015).

Other kinds of individual difference measures have yielded somewhat more robust associations with exploratory or exploitative behaviors. Experiences of lifetime scarcity were related to decreased resource-maximizing decision-making (Chang, Jara-Ettinger, and Baskin-Sommers 2022) and individuals with adverse childhood experiences explored less in a foraging task (Lloyd, McKay, and Furl 2022). Contextual effects in foraging tasks affect the explore-exploit tradeoff, with greater acute stress yielding overexploitation (Lenow et al. 2017), increased arousal associated with increased levels of exploration, and increases in valence substantially increased exploitation (van Dooren et al. 2021). Further, there are associations between psychopathologies and explore-exploit decisions. Some examples include that smokers make less initial exploratory choices (Merideth A. Addicott et al. 2012), people with greater anxiety and depression use lower levels of directed exploration (R. Smith et al. 2021), subjects with alcohol use disorders or binge eating disorders showed decreased exploration when confronted with losses (L. S. Morris et al. 2016), and people with schizophrenia overuse random exploration strategies (Speers and Bilkey 2023). Taken together, explore-exploit tasks have been applied in a variety of psychological domains, yielding little consistency in terms of economic decisions, though people with maladaptive psychological or psychiatric attributes have an attenuated ability to optimize these decisions.

### 1.2 Neurobiological Mechanisms of Exploration and Exploitation

Explore-exploit tasks lack behavioral convergence, contain a multitude of possible independent variables, and lack a coherent theory as to whether exploration and exploitation are products of disparate versus unified mechanisms. Given the lack of clarity regarding the constructs and behaviors guiding explore-exploit decisions, another approach could examine the neurobiological factors that are consistent across explore-exploit choices. One notable challenge that could be observed neurobiologically is if explore-exploit tasks elicit a consistent or disparate set of responses during exploration versus exploitation (Cohen, McClure, and Yu 2007) *(see Figure 1).* If exploration and exploitation elicit reliably different patterns of activation across various tasks, it could provide a window into the mechanisms may modulate explore-exploit decisions through an opponent processing system. Over the past two decades, the accumulation of neuroimaging studies conducted in explore-exploit tasks suggests that reviewing these common patterns may provide insight into explore-exploit decision making as a whole.

**Figure 1:**
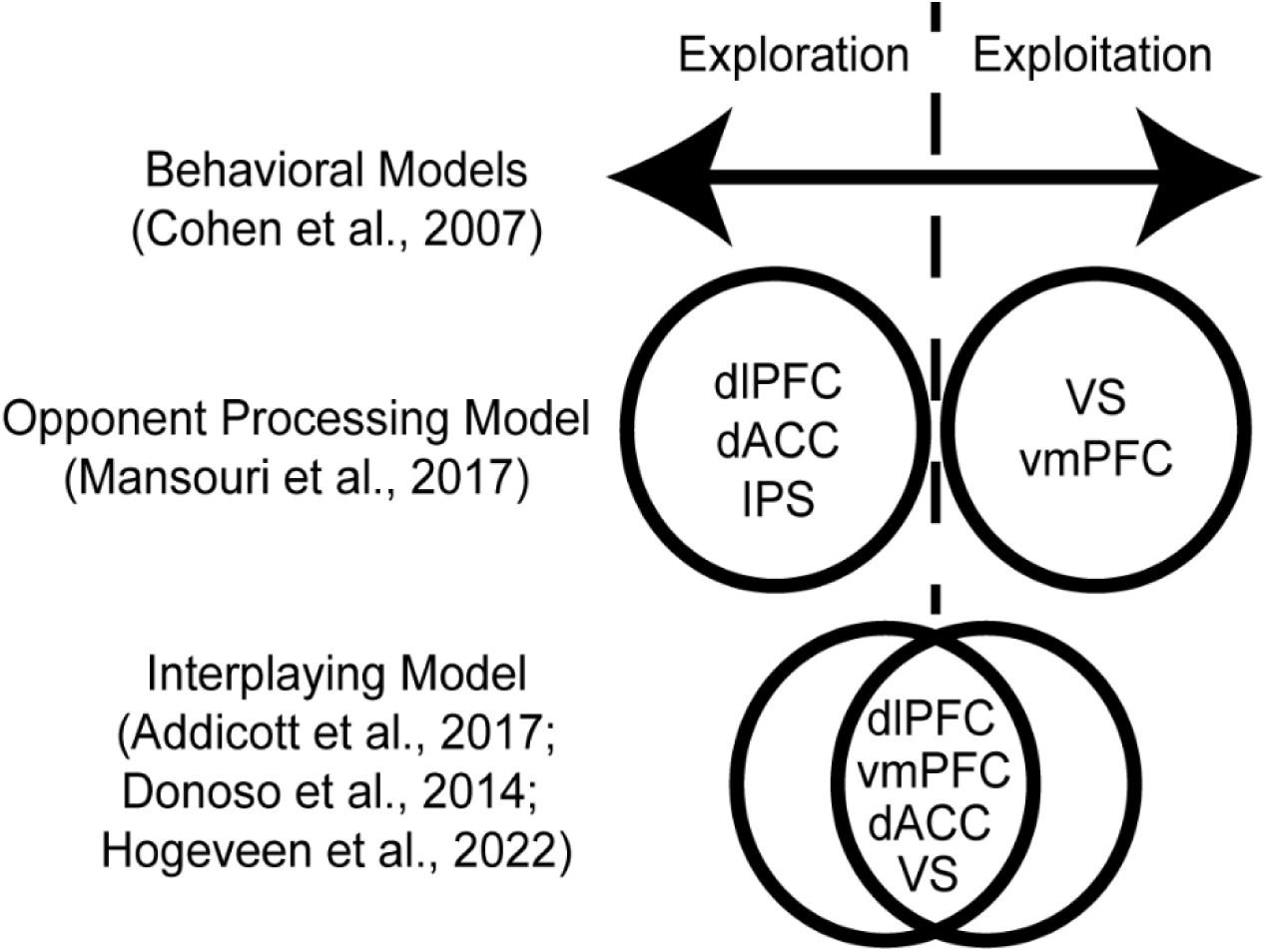
Models of exploration and exploitation. Behavioral models of explore-exploit decisions envision an optimal set of decisions, with people expressing tendencies to over or underexploit on a continuum of possible options. Nonetheless, the underlying mechanisms may be a binary or opponent processing model, where people are either exploring or exploiting, with corresponding brain activation. Another model suggests that exploration and exploitation is the product of the interplaying of underlying variables, where all the brain regions are involved in both exploration and exploitation, though may work together more or less depending on the explore-exploit situation.

Explore-exploit decisions in neuroscience have identified several key cortical and subcortical brain regions that contribute to these choices (Dennison, Sazhin, and Smith 2022). In animal literature, the Anterior Cingulate Cortex (ACC) has been identified as a major modulator of explore-exploit decisions. Versions of the n-armed bandit task have been adapted for rats and mice using n-armed radial mazes (Ohta et al. 2021). ACC activation has been linked to foraging in rats through an adapted patch foraging task (Kane et al. 2022) and a two-armed bandit monkey lesion study (Kennerley et al. 2006). Similarly, the dACC is tied to both exploration and exploitation in a monkey foraging task (Ramakrishnan, Hayden, and Platt 2019; Hayden, Pearson, and Platt 2011). Nonetheless, other findings suggest that the ventral striatum (VS) and amygdala represent immediate and future value of exploratory choices respectively in rhesus monkeys (Costa, Mitz, and Averbeck 2019). In human neuroimaging studies, there are some commonly cited areas of activation in brain regions associated with cognitive control (dlPFC), reward (VS), and attention (ACC), which are often used in region of interest (ROI) analyses (Shenhav et al. 2014; Wiehler, Chakroun, and Peters 2021; Hogeveen et al. 2022).

### 1.3 Evidence for Models of Exploration and Exploitation

While it is known that an array of brain regions is involved in explore-exploit decisions, it remains a key challenge to understand how these brain regions respond to dynamic environments. Two major accounts explain explore–exploit behaviors, which are the interaction of several neural regions depending on contextual features of the explore-exploit decision (e.g., dACC, dorsal striatum, lateral PFC, and VS; Donoso et al., 2014), or a dual-system driven by opponent processes of exploration and exploitation in frontoparietal regions (e.g., dlPFC, **d**ACC, IPS vmPFC and VS; Mansouri et al. 2017; Hogeveen et al. 2022) (*see Figure 1)*. If exploration and exploitation are generally more context-dependent rather than dissociable constructs, there may be less consistency in activation across these decision phases and across tasks. Instead, clusters of brain regions may work together depending on the task or context resulting in exploration or exploitation behavior (Addicott et al., 2017, Donoso et al., 2014, Hogeveen et al., 2022). With this interpretation, exploration and exploitation behaviors may be more context specific, and could potentially better described by underlying variables such as the risk, uncertainty, information, time horizons or other variables involved in the decision-making process. Thus, an interplayed model may be the combination of underlying psychological constructs in a given situation. Nonetheless, one challenge of an interplayed model is that there could also be a more complex set of responses to exploration or exploitation, potentially represented through connectivity patterns in the brain. It is also plausible that exploration and exploitation can be construed as opponent processes with concrete neural markers that reliably switch between exploration and exploitation (Mansouri et al., 2017). If this model has greater support, exploration and exploitation would be dissociable across tasks with consistent neural markers of activation. Furthermore, if there is evidence of opponent processing in exploration and exploitation this could allow for targeted interventions aimed at modulating these behaviors.

In trying to reconcile these accounts, studies tease apart how certain elements of explore-exploit dilemmas contribute to those decisions. For instance, understanding how environmental uncertainty and trends in information mediate this process may inform some of the underlying mechanisms in explore-exploit choices, with environmental uncertainty of new options (Badre et al. 2012; Navarro, Newell, and Schulze 2016; Tomov et al. 2020) seemingly largely processed in the PFC. Uncertainty in an environment has been represented in the brain in several ways, with relative uncertainty in the right rostrolateral PFC (Badre et al. 2012) and striatal dopamine function (Frank et al. 2009) driving directed exploration. The vmPFC was implicated in representing environmental uncertainty (Trudel et al. 2020), evidence accumulation in switching decisions (Blanchard and Gershman 2018), and determining the value of well-defined foraging options (Kolling et al., 2012). Taken together, these findings reinforce the importance of both frontopolar and subcortical regions in explore-exploit decisions, though it remains unclear to what degree an opponent process model driven by the frontoparietal cortex is supported by the weight of the evidence.

In sum, the current state of knowledge is limited in identifying consistent elements supporting neural circuitry associated with explore-exploit decisions, whether there are systematic biases in the literature, and if certain brain regions remain underemphasized in the reporting and interpretation of the data. One means of addressing these limitations is through quantitatively assessing patterns of activation across neuroimaging studies using coordinate-based meta-analyses (CBMA). We hypothesized that there would be convergence across explore-exploit studies in the activation of the vmPFC, dlPFC, VS, ACC, and IPS during explore-exploit decisions. These decisions are differentiated from the feedback phase where participants receive rewards based on their decision. While the feedback phase can provide important information for encoding the value of current and alternative outcomes while receiving feedback (Boorman, Rushworth, and Behrens 2013; Tsujimoto, Genovesio, and Wise 2011) the feedback phase does not completely capture the decision to shift choices on the following turn. Next, we expected that the exploitation versus exploration decision would be associated with greater vmPFC, VS, ACC activity in the exploration phase and that the exploration phase would be associated with greater activation in the IPS and dlPFC than in the exploitation phase. We speculated that the ACC could reflect an important role during exploitation through conflict monitoring (Weissman et al. 2003; Kim, Kroger, and Kim 2010; Mansouri et al. 2017) as exploitation involves decisions when to switch back to exploration. Thus, we expected that exploitation may reflect other processes beyond simply reward consumption since there is a routine opportunity to revert back to exploration. The act of monitoring this tradeoff and avoid over-exploiting could be a result of ACC conflict monitoring.

Since we began this investigation, two groups of researchers have conducted meta-analyses of explore decisions (Zhen et al. 2022), finding that exploration results in consistent activation of the dorsal medial prefrontal cortex and anterior insula, dorsolateral prefrontal cortex inferior frontal gyrus, and motor processing regions. (Wyatt et al. 2024). We extend these results by comparing exploration versus exploitation with a larger sample of studies and investigating task-based differences between exploration and exploitation using Seed-based D Mapping (SDM) software. Including the contrast between exploration and exploitation serves as a crucial means to subtract the effects of value-based decision making in order to understand activation that is unique to exploration and exploitation. Another group argued that prefrontal and parietal circuits integrate and switch between exploration and exploitation. Our approach differs from (Wyatt et al. 2024) in that we conducted a quantitative rather than qualitative meta-analysis, with regions identified subsequent to conservative thresholding and permutation testing. Overall, our results aims to identify activation patterns that are unique to exploration and exploitation, thereby helping identify to what degree we can theoretically understand these choices within an opponent processing model. We also explore activation differences between n-armed bandits and other types of explore-exploit tasks while making exploration or exploitation decisions. In summary, we investigate the common patterns of activation across explore-exploit tasks, whether there are systematic biases in the literature, and if there are other regions that are underemphasized in the interpretation of the data.

## 2. Materials and Methods

### 2.1 Inclusion Criteria and Study Selection

The current coordinate-based meta-analysis primarily followed PRISMA guidelines for meta-analyses regarding inclusion criteria, filtering, and analyses (Moher et al. 2009). We incorporated a pre-registration (https://aspredicted.org/7hc7c.pdf), which detailed the hypotheses and analyses we intended to use. We conducted a systematic literature search to identify explore-exploit studies that used neuroimaging techniques. First, we identified search terms by examining task names from several existing explore-exploit literature reviews (Cohen, McClure, and Yu 2007; M. A. Addicott et al. 2017; Zhen et al. 2022). Potentially eligible studies published through 1/01/2023 were identified by searching the PUBMED using the grouped terms: (n-armed OR exploration-exploitation OR explore-exploit OR multi-armed OR forage OR foraging OR “reward rate” OR (explore AND exploit) OR “reward trend” OR “clock task” OR clock-task OR “temporal-difference” OR “patch leaving” OR patch-leaving OR leave-stay OR “time horizon” OR “horizon task” OR bandit OR MVT OR “marginal value theorem” OR leapfrog OR “leap frog” OR leap-frog OR prey model OR “diet breadth model” OR “web surfing task” OR “web-surfing task” OR trend-guided OR “uncertainty driven”) AND (fMRI OR “functional magnetic resonance imaging” OR neuroimaging OR brain OR neural OR MNI OR “Montreal Neurological Institute” OR Tal OR coordinates). To enhance search sensitivity, the reference lists of the retrieved articles and review papers were further checked to identify potentially relevant articles. Additionally, we included studies that reported whole-brain analyses, as region of interest based analyses can bias coordinate-based meta-analyses (Moher et al. 2009) and were thus excluded. Finally, we incorporated studies that reported coordinates in a standard stereotactic space [i.e., Talairach or Montreal Neurological Institute (MNI) space]. The search process was conducted by Avi Dachs, with the first author identifying the studies accepted for final inclusion in the meta-analysis. For eligible studies that did not report whole-brain data, we contacted authors if the required information was unavailable in the published reports.

The initial PUBMED search yielded 6,214 papers. Of these, 5,256 papers were then excluded based on title, leaving 958 papers to be excluded by abstract and full text contents. Of the 958 remaining papers, 762 papers were excluded for not covering explore and exploit tasks, 72 relevant papers were excluded for not collecting fMRI data, 45 animal studies were excluded, and 14 non-empirical papers were excluded, leaving only 65 papers for data extraction and coding (see Figure 2). In the coding phase, 47 more papers were excluded due to data that were incompatible with our analysis (i.e., not fMRI or whole-brain), leaving a total yield of 19 papers. Finally, our list of papers was cross-referenced with the papers included in a similar meta-analysis (Zhen, 2022) revealing 4 papers that had been wrongly excluded from our search. After these papers were added, our final corpus included 23 papers with a cumulative N of 602 participants (see Figure 2 and Table 1). In total, we included 13 n-armed bandit studies, which varied in the number of bandits presented to the participant. We identified foraging tasks and 3 other tasks, including a problem-solving task, clock hand task, web surf task, and an observe-bet task. We grouped non-n-armed bandit tasks into an “other” category with a total of 10 studies to serve as a comparison group. Unlike n-armed bandits, the “other” tasks do not employ feedback about exploration or exploitation on each turn. Foraging, web-surf, and observe-or-bet tasks have clear shifts between exploration and exploitation based on observable changes in strategy. The clock hand task employs a fixed reward structure which is learned over time and exploration and exploitation is classified based on response times (M. A. Addicott et al. 2017). Thus, we classify tasks that do not have a continuous sequence of changing rewards as “other” types of exploration and exploitation tasks. Further, n-armed bandits involve *inferred* shifts to exploitation, whereas foraging tasks have distinct shifts from exploiting to traveling to other patches (Barack 2024). While both n-armed bandit and foraging tasks are grouped as explore-exploit tasks, they are sufficiently different to serve as potential comparison groups.

**Figure 2:**
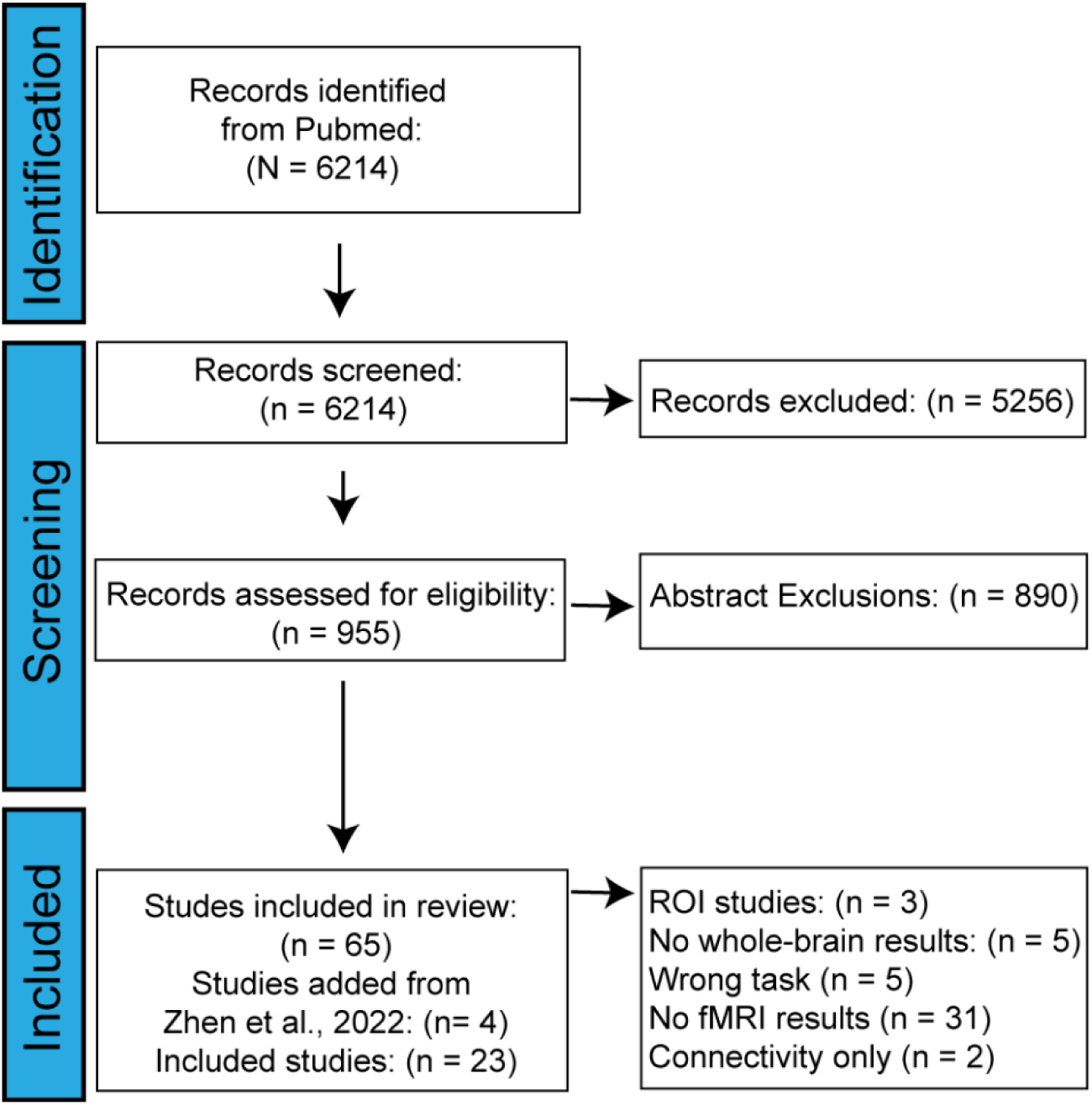
Inclusion criteria. Using the PRISMA flowchart system, we report the studies we identified, screened, and included in the meta-analysis. After we completed our initial search, four more studies identified by another meta-analysis (Zhen et al., 2022) that assessed activation during exploration. Starting from an initial search of 6,214 studies, we included n = 23 studies in our analyses.

**Table 1:** Included studies. N = 23 studies were included for the meta-analysis during explore-exploit decisions, and for the study of activation during exploration and exploitation specifically.

| Paper | Program | Threshold | N | Age | Females | Task |
| --- | --- | --- | --- | --- | --- | --- |
| Aberg et al., 2022 | SPM | 0.001 | 28 | 27.6 | 20 | 3-armed bandit |
| Abram et al., 2019 | FSL | 0.05 | 25 | 28.0 | 12 | web-surf task |
| Addicott et al., 2014 | SPM | 0.005 | 22 | 36.0 | NA | 6-armed bandit |
| Amiez et al., 2012 | AFNI | 0.05 | 11 | 27.9 | 6 | 4-armed bandit |
| Badre et al., 2012 | SPM | 0.001 | 15 | 20.0 | 8 | clock hand task |
| Blanchard & Gershman, 2018 | SPM | 0.001 | 18 | 26.0 | 11 | observe-bet |
| Chakroun et al., 2020 | SPM | 0.05 | 31 | 26.9 | 0 | 4-armed bandit |
| Daw et al., 2006 | SPM | 0.001 | 14 | NA | NA | 4-armed bandit |
| Donoso et al., 2014 | SPM | 0.005 | 40 | 18-26 | 20 | 4-armed bandit |
| Dunne et al., 2016 | SPM | 0.005 | 17 | 23.3 | 9 | 2-armed bandit |
| Howard-Jones et al., 2020 | SPM | 0.001 | 16 | 25.5 | 8 | 4-armed bandit |
| Kolling et al., 2012 | FSL | 0.05 | 20 | 22-32 | 12 | foraging |
| Korn et al., 2018 | SPM | 0.001 | 28 | 23.5 | 13 | foraging |
| Korn et al., 2019 | SPM | 0.001 | 24 | 25.0 | 11 | foraging |
| Laurerio-Martinez et al., 2014 | SPM | 0.05 | 50 | 35.1 | 11 | 4-armed bandit |
| Laurerio-Martinez et al., 2015 | SPM | 0.05 | 63 | 35.2 | 11 | 4-armed bandit |
| Mobbs et al., 2013 | SPM | 0.05 | 15 | 25.0 | 10 | foraging |
| Seymour et al., 2012 | SPM | 0.001 | 30 | NA | NA | 4-armed bandit |
| Tomov et al., 2020 | SPM | 0.001 | 31 | 18–35 | 17 | 2-armed bandit |
| Trudel et al., 2020 | FSL | 0.05 | 24 | 25.6 | 14 | 2-armed bandit |
| Wang and Voss., 2014 | AFNI | 0.05 | 42 | 18-35 | 22 | foraging |
| Wittman et al., 2016 | FSL | 0.05 | 20 | 21–32 | 8 | foraging |
| Zacharopoulos 2018 | SPM | 0.001 | 18 | 18-37 | 11 | foraging |

### 2.2 Statistical Analysis

We conducted a CBMA meta-analysis using Seed-based d mapping software (SDM-PSI version 6.22). SDM was implemented using several steps to conduct analyses described in the SDM tutorial and manual. First, we imported the screened data by preparing folders with MNI text files that reported the clusters and t values for each coordinate. Files were named in accordance with SDM guidelines, which requires specifying if the coordinates were derived from FSL, SPM, or “other” software, and whether the coordinates were in MNI or Talairach space.

For studies that reported Talairach coordinates, these results were automatically converted Talairach inputs into MNI space for subsequent analyses. Exploration and exploitation decisions were grouped based on the constructs reported in each study. 11 studies reported explore>exploit and exploit>explore contrasts and were coded as exploration and exploitation respectively (see Table 2). Other studies reported parametric effects for exploration and exploitation decisions (see Table 2) through assessing several components underlying value-based decisions in uncertain environments. Studies modulated uncertainty (Badre et al. 2012; Trudel et al. 2020), relative value (Howard-Jones et al. 2010), task difficulty (Abram et al. 2019), search evidence, and search cost across decision stages (i.e.., exploration in Stage 1 and exploitation in Stage 2) (Zacharopoulos et al. 2018). We classified reinforcement learning associated with recent experience as exploration (Dunne, D’Souza, and O’Doherty 2016).

**Table 2:** Reported contrasts. Some studies reported explore>exploit and exploit>explore contrasts, which were coded as exploration and exploitation respectively. Others reported parametrically modulated effects. The classification of those modulators as exploration and exploitation is described in the methods section. The t-threshold identified for each study is reported, which was determined by the threshold identified in the analyses conducted by each study and their respective sample size.

| Paper | T -threshold | Contrasts |
| --- | --- | --- |
| Aberg et al., 2022 | 3.6459 | Explore>Exploit |
| Abram et al., 2019 | 2.0639 | Parametric Modulator |
| Addicott et al., 2014 | 3.1352 | Explore>Exploit |
| Amiez et al., 2012 | 2.2282 | Explore>Exploit |
| Badre et al., 2012 | 4.1404 | Parametric Modulator |
| Blanchard & Gershman, 2018 | 3.9216 | Explore>Exploit |
| Chakroun et al., 2020 | 2.0423 | Explore>Exploit |
| Daw et al., 2006 | 4.2208 | Explore>Exploit |
| Donoso et al., 2014 | 2.0227 | Parametric Modulator |
| Dunne et al., 2016 | 3.252 | Parametric Modulator |
| Howard-Jones et al., 2020 | 4.0728 | Parametric Modulator |
| Kolling et al., 2012 | 2.093 | Parametric Modulator |
| Korn et al., 2018 | 3.6896 | Explore>Exploit |
| Korn et al., 2019 | 3.7921 | Explore>Exploit |
| Laurerio-Martinez et al., 2014 | 2.0096 | Explore>Exploit |
| Laurerio-Martinez et al., 2015 | 1.999 | Explore>Exploit |
| Mobbs et al., 2013 | 2.1448 | Parametric Modulator |
| Seymour et al., 2012 | 3.6594 | Explore>Exploit |
| Tomov et al., 2020 | 3.6459 | Parametric Modulator |
| Trudel et al., 2020 | 2.0686 | Parametric Modulator |
| Wang and Voss 2014 | 1.68 | Parametric Modulator |
| Wittman et al., 2016 | 2.093 | Parametric Modulator |
| Zacharopoulos 2018 | 3.9651 | Parametric Modulator |

Another variation of reinforcement learning included assessing exploitation as the last average reward rate, whereas exploration reflecting the expected values associated with learning past reward rates (Wittmann et al. 2016). Another study coded foraging value-decision value contrast for exploration and search value-decision value contrast for exploration (Kolling et al. 2012).

Another study varied the parametric value of staying with a foraging patch whereas exploration was classified as the difference in decision versus consumption (Abram et al. 2019). Switch-in events were classified as exploration and activation associated with actor absolute reliability as exploitation due to the behavioral design (Donoso, Collins, and Koechlin 2014). Other studies varied the presence or absence of newly available information (Wang and Voss 2014), and the advantageousness of the environment (Mobbs et al. 2013). In another study, the modeled choice kernel reflected exploratory decisions (Seymour et al. 2012). These classifications of exploration and exploitation reflect the coordinates selected for analysis and are accessible on OSF. Overall, the parametric effects generally reflected sensitivity to value, information, or uncertainty while exploring or exploiting an uncertain environment.

Next, we created an SDM table with all the respective peak coordinates. We noted t-stats in the SDM table with respect to effect sizes and converted reported p and z stats using the SDM “convert peaks” function (see Table 2). Then, we completed preprocessing using Functional MRI as its modality, with a gray matter correlation matter template following validated methods (Albajes-Eizagirre et al. 2019; J. Radua et al. 2012). We used a 1.0 anistropy setting, a 20 mm FWHM isotropic kernel, a gray matter mask, and a standard 2mm voxel size. This was followed by a mean analyses with 50 imputations (Joaquim Radua and Mataix-Cols 2009). To compare exploration and exploitation decisions, and n-armed bandit versus other tasks, we generated linear models respectively where we compared these groups by assigning a linear model analysis (Joaquim Radua et al. 2010). We used the SDM meta-regression tool with prediction dummy variable {exploit=1, explore=0} and {n-armed=1, other=0} for the positive side of the significance test. These models thus contrasted exploit with explore conditions and n-armed bandit with other tasks respectively, with a 1 selected for the contrast of interest in the SDM model. Additionally, we included several nuisance regressors to control potentially confounding variables, specified as a “0” in the SDM linear model. Specifically, we included analysis type (parametrically modulated = 1, unmodulated = 0) and the smoothing kernel size (Sacchet and Knutson 2013). We included these nuisance regressors to ensure that analysis type was not a confounding variable and since the size of the smoothing kernel can move the observed activation anterior or posterior of the brain.

Subsequently, we performed family wise error corrections and using n=1000 permutations (Albajes-Eizagirre et al. 2019). This correction controls for multiple comparisons by randomly swapping the effect-sizes between the voxels for each study, recalculating the means of the studies for each voxel and saving the maximum of the means (Albajes-Eizagirre et al. 2019). The results were then thresholded using threshold-free cluster enhancement (TFCE) with a corrected probability threshold of *p* < 0.05 (S. M. Smith and Nichols 2009). TFCE has also been shown to have a sensitivity comparable to a Family Wise Error correction and yield valid results with only about five percent of significant clusters based on spurious convergence across 200,000 simulated meta-analyses (Frahm et al. 2022). TFCE statistics are generally neither too liberal or conservative and have been used across many meta-analyses (Albajes-Eizagirre et al. 2019; Masson et al. 2021; Sheng et al. 2020). Masks were created and their values were extracted for reporting. For the conjunction of explore and exploit conditions, we conducted a CBMA of explore and exploit conditions respectively, and then used the multimodal function provided by SDM to produce the conjunction map.

To assess potential heterogeneity and potential bias in the CBMA results, we extracted funnel plots. We report the strength of evidence through multiple robustness considerations, study heterogeneity (I² statistic), effect of small studies on the results (metabias) with resulting funnel plot asymmetry, and excess significance. The funnel plots are constructed through assessing the residual, or the weight each study has in the meta-analysis, with the size of its treatment effect, identified as precision on the y axis, though these tests must be interpreted with caution as publication bias can arise from multiple sources (Sterne et al. 2011). All analyses were completed in Montreal Neurological Institute (MNI) space. To report consistent results across human brains (Mazziotta et al. 1995), we show probabilistic anatomical labels for clusters of activation using the Harvard–Oxford cortical and subcortical atlases (Desikan et al. 2006).

## 3. Results

We completed meta-analyses that assessed activation across explore-exploit tasks, followed by activation specific to exploration and exploitation. All meta-analyses controlled the size of the smoothing kernel, as well as whether the analyses reported were parametrically modulated or unmodulated. The first meta-analysis investigated activation pooling across both exploration and exploitation conditions and is reported in *Supplementary Results.* Next, we assessed the contrast between exploration and exploitation. Subsequently, we report activation that is consistent across both exploration and exploitation. Lastly, we show exploratory results revealing activation that is greater among n-armed bandit tasks versus other tasks during exploitation and exploration.

### 3.1 Neural Responses Between Exploration versus Exploitation Phases

We conducted a CBMA contrasting the exploration and exploitation conditions across all the explore-exploit tasks. We hypothesized that the exploitation phase would be associated with greater vmPFC, VS, ACC activity than the exploration phase. We did not find any significant clusters for exploitation versus exploration that exceed a threshold of *p* < .05. Next, we hypothesized that the exploration phase would be associated with greater activation in the IPS and dlPFC than in the exploitation phase. Our results indicated five significant clusters of activation in the dorsolateral prefrontal cortex, right dorsal anterior cingulate cortex, anterior insula, and superior temporal gyrus *(see Figure 3, Table 3)*. Using the Harvard-Oxford Atlas, our results were consistent with our hypotheses in finding stronger activation in the dlPFC during exploration versus exploitation. NeuroSynth was used to confirm anatomical localization, in conjunction with direct anatomical references from prior literature. We verified our anatomical labels for the centroids of the clusters using NeuroSynth with the first cluster (24,2,60) being associated with the term “dorsal premotor” (z = 7.55), second cluster (10,22,40) associated with the term “anterior cingulate cortex” (z = 7.96) and “dacc” (z= 4.96), the third cluster (34,22,6) being associated with the term “anterior insula” (z = 10.39), the fourth cluster (58,0,0) term associated with “superior temporal” (z = 7.13) and the fifth cluster (32,36,26) associated with the term “dorsolateral” (z=4.19). We followed up with analyses of metabias and excess significance, finding no significant metabias or excess significance for this CBMA.

**Figure 3.**
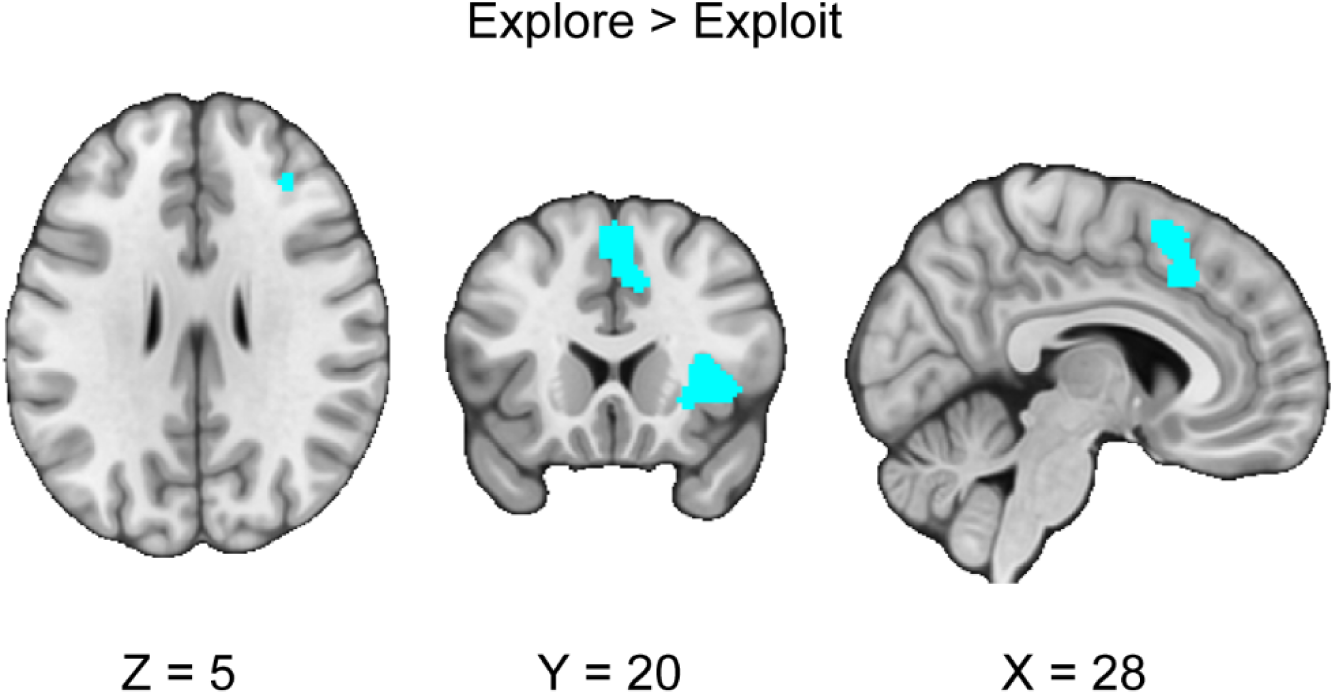
Clusters of activation in the contrast between explore versus exploit phases. Our results indicated that during the exploration versus exploitation phases, we found stronger activation in the dlPFC, AI and dACC, and the superior temporal gyrus. Maps were thresholded using TFCE at *p*=.05 and rendered in MRIcroGL. Thresholded (https://neurovault.org/images/888157/) and unthresholded (https://neurovault.org/images/888156/) images are available at Neurovault.

**Table 3:** Reported clusters of activation across meta-analyses conducted. Confirmatory analyses conducted according to the pre-registration include the explore and exploit conditions and assessing their respective conjunctions and contrasts. Exploratory analyses were conducted on n-armed bandit versus other tasks. Coordinates are reported in Montreal Neurological Institute (MNI) space.

| Analysis Type | MNI coordinate | SDM-Z | P | Voxels | Cluster Location |
| --- | --- | --- | --- | --- | --- |
| Confirmatory |  |  |  |  |  |
| Explore > Exploit | 24,2,60 | -4.784 | 0.001 | 1310 | Right superior frontal gyrus, dorsolateral, BA 6 |
|  | 10,22,40 | -3.03 | 0.012 | 838 | Right median cingulate / paracingulate gyri, BA 32 |
|  | 34,22,6 | -2.994 | 0.013 | 596 | Right insula, BA 48 |
|  | 58,0,0 | -2.882 | 0.24 | 266 | Right superior temporal gyrus, BA 48 |
|  | 32,36,26 | -4.205 | 0.02 | 41 | Right middle frontal gyrus, BA 46 |
| Exploit < Explore | Null |  |  |  |  |
| Exploratory |  |  |  |  |  |
| N-armed > Other (Exploit) | Null |  |  |  |  |
| Other > N-armed (Exploit) | 4,34,44 | -4.296 | 0.003 | 475 | Right superior frontal gyrus, medial, BA 8 |
|  | 30,24,4 | -4.468 | 0.003 | 352 | (undefined), BA 47 |
| N-armed > Other (Explore) | Null |  |  |  |  |
| Other > N-Armed (Explore) | Null |  |  |  |  |

### 3.2 Neural Responses to the Conjunction Between Exploration and Exploitation Phases

We conducted a conjunction analysis across exploration and exploitation in the sample of studies we collected. We hypothesized that the conjunction of explore and exploit phases would be associated with activation in the IPS, dACC, and dlPFC. Supporting our hypothesis, we found common activation in the dACC in the conjunction between exploration and exploitation decisions *(see Figure 4).* Activation in exploration and exploitation elicits common activation across multiple regions with a correlation of *r* = .62 across the unthresholded explore and exploit images. These results suggest that areas of common activation should in the future be closely examined using multivariate and connectivity methods to understand how they are involved in exploration and exploitation. In contrast to our hypothesis, we did not find convergence in the IPS or dlPFC. We also found conjunctive patterns of activation in the dmPFC and anterior insula.

**Figure 4.**
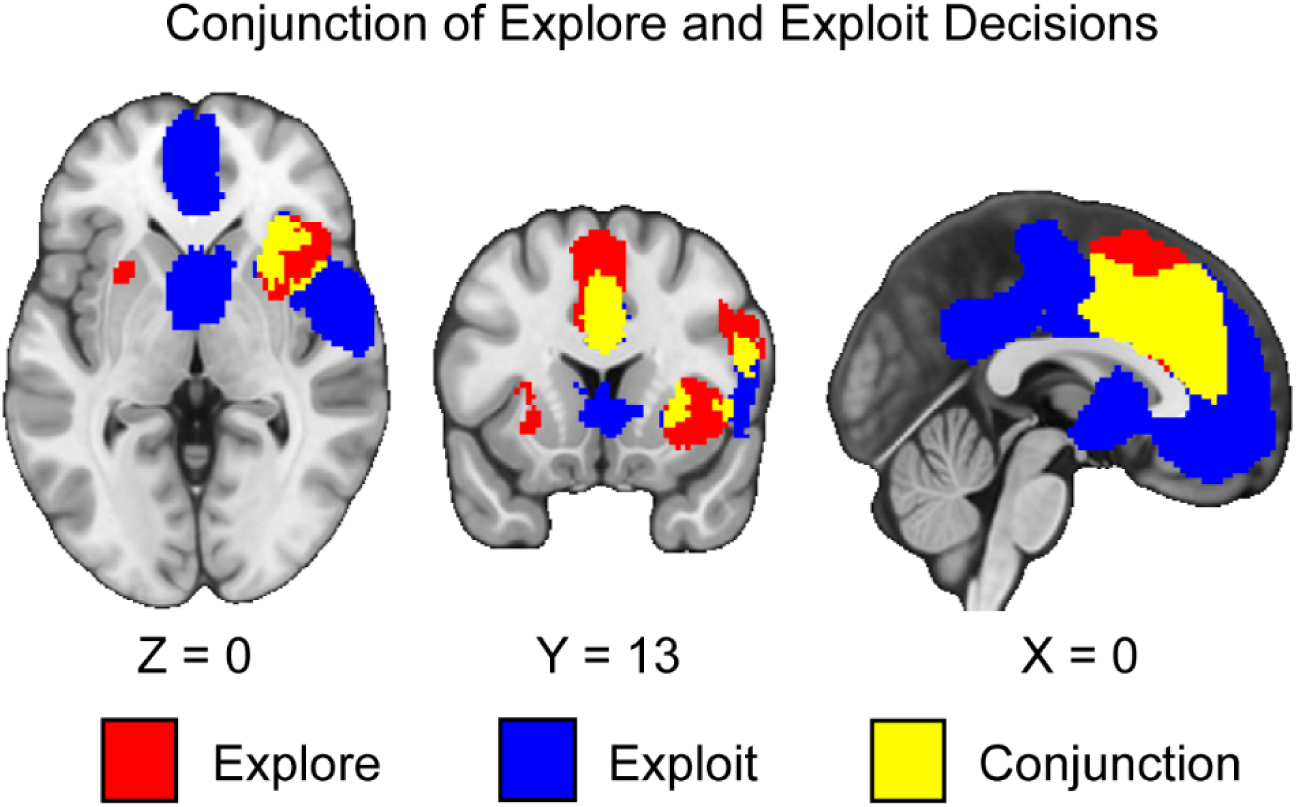
Clusters of activation in the conjunction of explore and exploit decisions. Our results indicate that during both explore and exploit phases, there is activation in the ACC, dmPFC, and the insula. Maps were thresholded using TFCE at *p*=.05 and rendered in MRIcroGL. Red corresponds to activation during exploration, blue during exploitation, and yellow for the conjunction between both decisions. Thresholded images are available at https://neurovault.org/images/888159/, https://neurovault.org/images/888160/, and https://neurovault.org/images/888161/. Unthresholded images are available at https://neurovault.org/images/888167/, https://neurovault.org/images/888168/, and https://neurovault.org/images/888169/.

### 3.3 Differential Activation Between N-Armed versus Other tasks During Exploration and Exploitation

We followed up our pre-registered hypotheses by assessing if there are differences between activation in n-armed bandit tasks compared to other tasks during exploration and exploitation. If there are activation differences between tasks, this may suggest that these tasks are not eliciting consistent patterns of activation in exploration and exploitation as may be expected.

During exploration, we did not find any activation differences between n-armed bandits and other tasks. During the exploitation phase, we found that other tasks versus n-armed bandits resulted in two significant clusters (*see Table 3, Figure 5*). There was no reported excess significance, or metabias in the results (*p* > .001).

**Figure 5.**
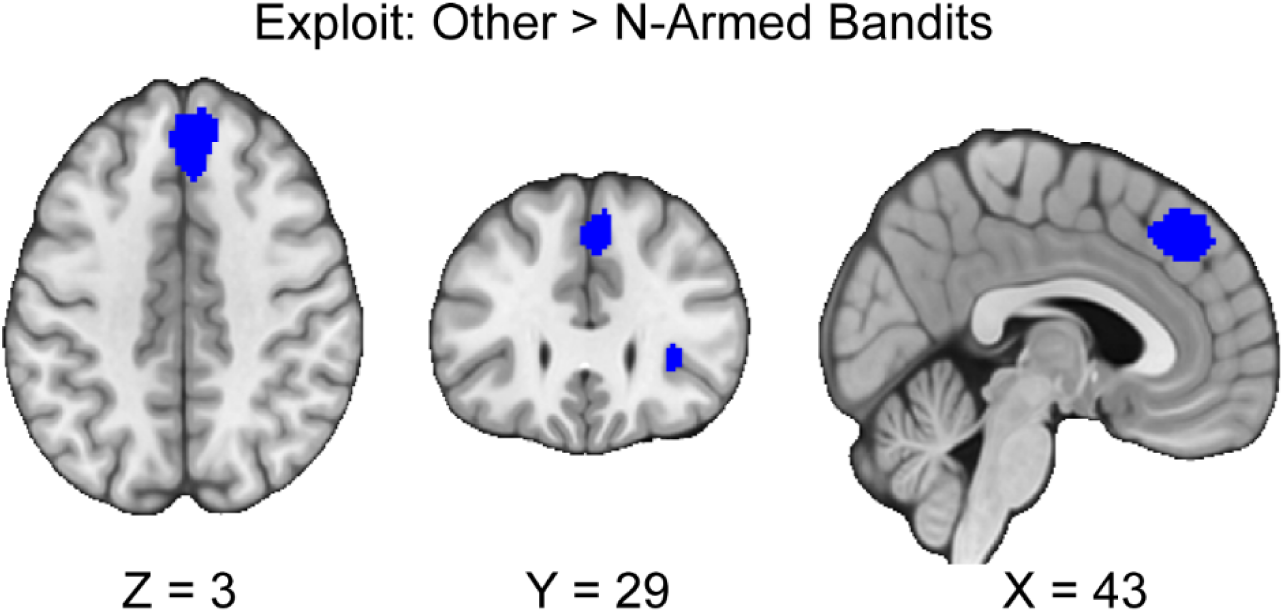
Clusters of activation during exploitation, between other tasks and n-armed bandits. Our results indicate that there are differences in activation between tasks during exploitation. We found that there was greater activation in the dmPFC and AI when participants exploited in other tasks versus n-armed bandits. Maps were thresholded using TFCE at *p*=.05 and rendered in MRIcroGL. Thresholded images are available at Neurovault: https://neurovault.org/images/888163/. Unthresholded images are available at Neurovault: https://neurovault.org/images/888162/, https://neurovault.org/images/888164/, https://neurovault.org/images/888165/, and https://neurovault.org/images/888166/.

## 4. Discussion

This investigation conducted a coordinate-based meta-analysis of explore-exploit tasks. We included both n-armed bandit and other types of explore-exploit tasks and analyzed them to assess patterns of activation that are consistent across explore-exploit decisions, as well as unique to exploration and exploitation decisions respectively, and differences in activation between tasks. First, we found consistent activation unique to exploration and exploitation decisions, with activation in the dlPFC, vmPFC, ACC, IPS, dmPFC, and VS. These results suggest that exploration and exploitation generally evoke activation associated with value-based decision-making (Bartra, McGuire, and Kable 2013; Rangel, Camerer, and Montague 2008), although it does not control for activation unique to these individual decision phases. This suggests that while these regions are commonly recruited, their precise role may differ between exploration and exploitation. Second, we found greater activation in the dlPFC, dACC, and the AI during exploration versus exploitation. Third, we conducted an exploratory analysis to assess differences between n-armed bandits and foraging tasks during exploration and exploitation respectively. We found differences in activation in the AI and dmPFC between other tasks and n-armed bandit tasks during exploitation. Overall, our meta-analytic results support previous findings that have identified critical regions involved in exploration and exploitation.

### 4.1 Opponent Processing versus Interplaying Models of Exploration and Exploitation

When we investigated the contrast between exploration and exploitation, we found stronger activation in the dlPFC, AI, and dACC during exploration compared to exploitation, suggesting that these regions are part of opponent processes in explore-exploit decisions. These results are consistent with past findings (Laureiro-Martínez et al. 2014), suggesting that the dlPFC may contribute to tracking the value of alternative choices (Laureiro-Martínez et al. 2015; Raja Beharelle et al. 2015), attending to risk (Obeso et al. 2021), tracking uncertainty (Chakroun et al. 2020; Tomov et al. 2020), and guiding directed exploration (Zajkowski, Kossut, and Wilson 2017). Additionally, as the dlPFC is implicated in cognitive control (Friedman and Robbins 2022), dlPFC activation may affect working memory as it relates to information gain and integrating recent rewards (Cogliati Dezza, Cleeremans, and Alexander 2019; Cogliati Dezza et al. 2017).

The AI subserves several notable computational mechanisms that are relevant for exploration and exploitation. Overall, the AI has been found to respond more strongly exploration versus exploitation (Kayser et al. 2016; Ohira et al. 2013; Tomov et al. 2020; Blanchard and Gershman 2018). Two recent accounts suggest that the AI could be processing risk (Zhen et al. 2022), or could serve as part of a broader salience network during exploration (Hogeveen et al. 2022).

While the AI serves an important role within valuation processing (Bartra, McGuire, and Kable 2013), other studies have indicated that the AI is stronger activated with the sudden introduction of reward structures rather than stable reward systems (Li et al. 2006). Thus, while the AI is involved in risk processes (Preuschoff et al., 2011; Smith et al., 2014), its role may involve orienting the dACC and dlPFC toward changes in valuations related to risk and uncertainty.

Contrary to our expectations, we found that the dACC was more activated during exploration versus exploitation. The dACC may contribute to exploration versus exploitation by tracking trends in foraging tasks (Wittmann et al. 2016; Kolling et al. 2012), feedback processing (Kolling et al., 2016) preparing movement away from disadvantageous foraging patches (Mobbs et al. 2013), with more self-focused individuals showing lower activity in dACC compared to individuals who were foraging for others (Zacharopoulos et al. 2018), and evaluating salient feedback for learning optimal strategies (Amiez et al. 2012). The interpretations emphasizing the role of the dACC in foraging may be confounded as one investigation found that dACC engagement was explained by choice difficulty, and not the value of foraging (Shenhav et al. 2014). This account is further complicated by that recent in rhesus monkeys suggests that choice difficulty does not influence local field potentials in the dACC (Goussi-Denjean et al. 2023).These mixed findings suggest that the role of the dACC is difficult to categorize precisely given its heterogeneity in function across many cognitive processes. We judge that our results are consistent with the prefrontal and parietal circuits integrating and switching between exploration and exploitation (Hogeveen et al. 2022; Wyatt et al. 2024). Integrating the roles of the dlPFC, AI and dACC in regulating exploration versus exploitation is also consistent with recent findings suggesting that these regions could be part of a circuit that modulates strategic decisions (Jahn et al. 2023). Overall, our results broadly support the account that greater dlPFC, AI, and dACC activation during exploration versus exploitation can be characterized through opponent processes, though there are challenges in inferring the precise functions of each region during explore-exploit decisions.

We further acknowledge that while the center of the second cluster described captures the dACC, it also extends into the midcingulate cortex, and supplementary motor areas (Wagera et al. 2016). While the spatial resolution of CBMAs using FWER correction is relatively low, the cluster may potentially be reflecting activation and resulting processes outside of the dACC. Some studies indicate that the MCC is involved in attention processing (Touroutoglou et al. 2020), feedback processing, pain, salience, action-reward association, premotor functions, and conflict monitoring (Procyk et al. 2016). Additionally, SMA or dmPFC function could facilitate state change decisions by tracking reward rate over time (Kolling and O’Reilly 2018), and tracking reward magnitudes (Vural, Katruss, and Soutschek 2024). Thus, while the second cluster is consistent with past studies that have indicated the role of the dACC during exploration versus exploitation, this cluster may also be capturing several additional processes involved in explore-exploit decisions.

In contrast to recent meta-analyses (Wyatt et al. 2024; Zhen et al. 2022), our results suggest that many brain regions involved in value-based decision making are coactivated across both exploration and exploitation rather than evoking distinct patterns of activation. For example, our results suggested that the dorsal medial prefrontal cortex and premotor cortex (Zhen et al., 2022) were involved in both exploration and exploitation. While these brain regions may be involved in exploration, by subtracting activation related to exploitation we show that these other brain regions may be simply involved in the overall value-based decision process (Rangel, Camerer, and Montague 2008) rather than being unique to exploration. Additionally, while the qualitative approach taken by Wyatt and colleagues indicated that the IPS and Precuneus have greater activation during exploration and exploitation, our quantitative analyses indicate that many of these regions fail to survive thresholding and are generally sensitive to both exploration and exploitation. Thus, while we agree with a recent empirical work that control and attention networks are involved in exploration (Hogeveen et al. 2022), our results suggest that more precisely that greater activation in the dACC, AI, and dlPFC differentiates exploration and exploitation decisions.

However, there remain two large issues in interpreting exploration and exploitation through the lens of the opponent process model. The first is that there are more similarities than differences in activation across our results. Even when controlling for the effects of exploration and exploitation decisions specifically, our conjunction analyses reveal that exploration and exploitation generally elicit similar patterns of activation, particularly in the dACC and dmPFC.

These results suggest that areas of common activation should closely examined on the future using multivariate and connectivity methods to understand how they are involved in exploration and exploitation. Extending a previous meta-analysis suggests that these regions are not unique to exploration (Zhen et al. 2022), but are also involved in exploitation. As a result, when differences are reported in these regions, they may be due to the interplaying of more complex underlying variables modulating these brain processes rather than a product of a general opponent processing system for exploration versus exploitation decisions.

Secondly, our exploratory analyses suggest that there remains substantial heterogeneity between tasks. This issue may speak to the lack of behavioral convergent validity between these tasks (von Helversen et al. 2018), which is to say that a participant exploiting in a foraging task does not predict how they will exploit in an n-armed bandit task. During exploitation, we found differences in activation in the insula and dmPFC between other tasks and n-armed bandits. In theory, we would not expect to see differences in activation if exploitation across tasks reliably elicit similar responses, we would not expect to see differences between these tasks. Nonetheless, the differences in AI and dmPFC could reflect differences in how people perceive risk and uncertainty (ie: Zhen et al. 2022) or salient features (ie: (Hogeveen et al. 2022) while exploiting in n-armed bandits versus foraging tasks. Thus, while our results suggest that while the dACC, AI and dlPFC differentiate exploration and exploitation, these constructs remain fragile to the context of the decision based on task, and that most of the activation associated with these decision processes is indistinguishable and is modulated based on context. As such, the interplaying model of exploration and exploitation is generally a better descriptor of these constructs, though the dlPFC, AI, and dACC can act as opponent processes between these types of decisions.

### 4.2 Limitations

Although our work has found that exploration and exploitation can be dissociated by dlPFC, AI, and dACC activation, we acknowledge our study has several notable limitations. First, while we included N=23 studies, this quantity is fairly low for a CBMA type meta-analysis, with a common benchmark suggesting a minimum of 17–20 Experiments (Yeung et al. 2019). However, the exploratory CBMA of n-armed bandits versus other tasks during exploration and exploitation contrasted 13 versus 10 studies. Since this sample size is below the benchmark, it should be considered exploratory. Nonetheless, there is a lack of clear guidance as to what constitutes acceptable sample sizes for SDM, as this highly depends on the effects measured, the number of participants, and whether thresholded images are included or not. Second, while we found substantial areas of coactivation between explore-exploit conditions, we cannot conclude that these areas are consistently involved with both types of decisions. For example, prior studies have shown that a region may appear to be involved in different processes despite having patterns of activation and connectivity profiles (Woo et al. 2014; D. V. Smith, Sip, and Delgado 2015). Further analyses could disentangle the involvement of these brain regions and show distinct connectivity with other brain regions to better understand their involvement in explore-exploit decisions. Third, we acknowledge that CBMA has relatively low spatial resolution, limiting precise anatomical interpretations (Radua et al. 2012).

Other limitations extend beyond meta-analytic methods when assessing exploration and exploitation more generally. Explore-exploit tasks limit the manner in which information is presented, and a latent variable that may bias switching decisions includes the trend in information. Some studies have started to explore the effects of trends in information (Wittmann et al. 2016; Kolling et al. 2012; Vestergaard and Schultz 2020), though it remains underexplored how these trends bias people to act too soon or too late. Further, brain connectivity (Friston 2011; Utevsky et al. 2017) may reveal patterns of explore-exploit decision making, yet few connectivity studies (S. Morris et al. 2015; Tardiff et al. 2021) have been completed in this domain. Since the default mode network (DMN) is implicated in executive function and cognitive control (Fox et al. 2005), and the executive control network (ECN) serves to rapidly instantiate new task states (Marek and Dosenbach 2018), both the DMN and ECN could interact to drive exploiting versus exploring decisions. Future studies may reconcile the gap that remains in understanding how explore-exploit decisions are associated with brain connectivity patterns.

While acknowledging limitations for generalizing both behavioral and neural results resulting from exploration and exploitation, the finding that the dlPFC, AI, and dACC reliably distinguish exploration and exploitation could inspire important future directions. First, a fruitful future direction includes modulating dlPFC responses, which are quite common in transcranial stimulation studies. Since there are many links between the dlPFC and psychopathology such as schizophrenia (Wu et al. 2017), anxiety (Balderston et al. 2020), and substance use (Goldstein and Volkow 2011), regulating dlPFC activation may reliably modulate explore-exploit decisions. Specific to substance use, while there has been extensive research into the neural mechanisms of addiction, it remains underexplored how individual differences in decision making serve as risk factors for increasing consumption of substances. Past investigations revealed that smokers explore less and learn faster (Merideth A. Addicott et al. 2012) and require greater cognitive control when exploring (Merideth A. Addicott et al. 2014). People with greater alcohol use tend to avoid uncertainty (L. S. Morris et al. 2016) and explore less. Brain responses may be modulated by substance use and mediated by social context (Sazhin et al. 2020). Sharing rewards with friends decreases connectivity between VS and dorsomedial prefrontal cortex (Wyngaarden et al. 2023), suggesting that social contexts are an important feature of understanding substance use decisions. Future investigations could also study the role of trends in decision making and assess whether substance users forecast future trends worse than non-substance users. Using explore-exploit dilemmas, researchers can assess how people make predictions, and whether substance users have an impaired cognitive ability to predict future outcomes.

## 5. Conclusion

In summary, we conducted a coordinate-based meta-analysis of neuroimaging studies using explore-exploit tasks. We found that areas associated with executive control (dlPFC), attention (IPS, dACC), and reward (VS) are reflected in exploration and exploitation decisions.

Exploration versus exploitation can be distinguished by greater activation in the dlPFC, AI, and dACC. Nonetheless, there remains substantial heterogeneity in brain responses due to task types, modulated by activation in the AI and the dmPFC while exploiting. Further, exploration and exploitation are associated with more similar than dissimilar patterns of activation in the AI, dmPFC, dACC, and VS. These results suggest that exploration and exploitation are not reliable opponent processes but are more of a product of the interplaying of underlying physiological and psychological features guiding these decisions. Nonetheless, the finding that the dlPFC, AI, and dACC distinguish exploration and exploitation could serve as an important area of future research in cognitive neuroscience and psychopathology, as modulating these brain regions could shift how people explore and exploit.

## Supporting information

Supplemental Methods and Results

## Acknowledgments

This work was supported in part by a grant from the National Institute on Drug Abuse (R03-DA046733). We thank Chelsea Helion, Vishnu Murty, and Michael McCloskey for feedback on prior versions of this manuscript. We thank Amanda Nguyen for help with data collection.

## Data availability

Data is available on OSF at: https://osf.io/86kp9/. Thresholded and unthresholded statistical maps are located on https://neurovault.org/collections/17882/.

## Author Contribution Statement

Daniel Sazhin formulated the pre-registration, conducted analyses, and prepared the manuscript with contributions from all coauthors. Abraham Dachs collected the data and contributed to the manuscript. David Smith oversaw the project and contributed to the manuscript.

## Abbreviations

fMRI: Functional Magnetic Resonance Imaging
dlPFC: Dorsal Lateral Prefrontal Cortex
AI: Anterior Insula
ACC: Anterior Cingulate Cortex
dACC: Dorsal Anterior Cingulate Cortex
MCC: Midcingulate Cortex
IPS: Intraparietal Sulcus
vmPFC: Ventromedial Prefrontal Cortex
dmPFC: Dorsomedial Prefrontal Cortex
VS: Ventral Striatum

