## Supplemental Methods and Results for "Meta-Analysis Reveals That Explore-Exploit Decisions are Dissociable by Activation in the Dorsal Lateral Prefrontal Cortex, Anterior Insula, and the Dorsal Anterior Cingulate Cortex"

### **Supplemental Results**

Our first meta-analysis investigated activation across both exploration and exploitation. Exploration and exploitation were isolated through a contrast (ie. explore>exploit) or through the parametric effect of exploration or exploitation respectively. Additionally, this analysis controlled for effects of smoothing kernel size or analysis type (modulated or unmodulated). This omnibus analysis would reveal common patterns of activation specific to making decisions in explore-exploit situations. We hypothesized that there will be convergence across explore-exploit studies in activation in the vmPFC, dlPFC, VS, ACC, and IPS. Our results indicate one significant cluster of activation in the left superior frontal gyrus (see Supplemental Figure 1, Supplemental Table 1). Using the Harvard-Oxford Atlas, these clusters encompass the NAcc, paracingulate gyrus, and the Angular gyrus extending into the Supramarginal gyrus, providing meta-analytic evidence supporting these regions as centers of activation in explore-exploit situations. We followed up with analyses of metabias and excess significance, that none of the three clusters had excess significance or metabias (P<.001) given the expected given the power of the tests (Ioannidis and Trikalinos 2007). Overall, these results suggest that exploration and exploitation generally evoke activation in areas putatively associated with value-based decision making.

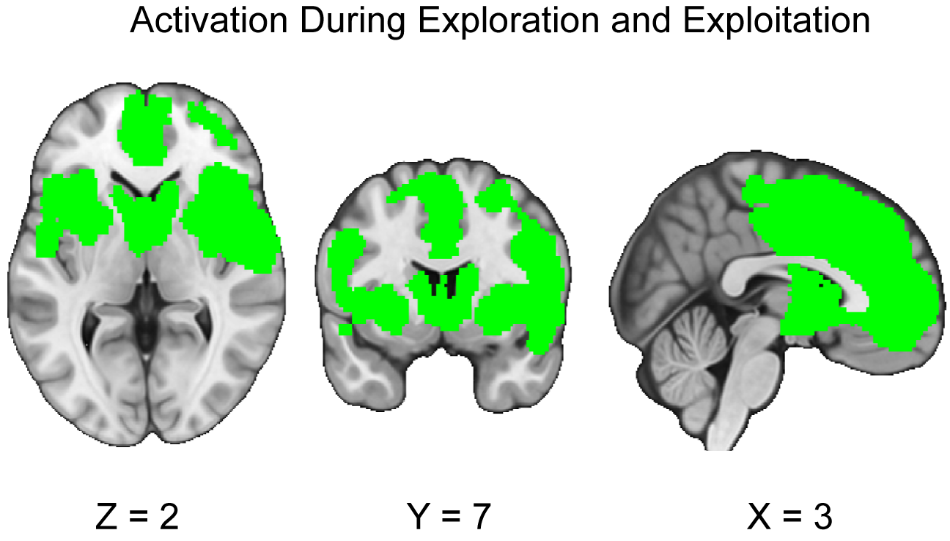

**Supplemental Figure 1:** **Clusters of activation (green) across all explore-exploit tasks, in both the exploration and exploitation conditions.** Our results are consistent with our hypotheses, showing activation in the ACC, IPS, dlPFC, vmPFC, and VS during exploration and exploitation. Maps were thresholded using TFCE at P=.05 and rendered in MRIcroGL. Thresholded and unthresholded images are available at: <https://neurovault.org/images/888154/> and <https://neurovault.org/images/888155/>.

| **Analysis Type** | **MNI coordinate** | **SDM-Z** | **P** | **Voxels** | **Cluster Location** |
| --- | --- | --- | --- | --- | --- |
| All | -4,3-,46 | 6.238 | 0.001 | 33274 | Left superior frontal gyrus, medial |

**Supplemental Table 1:** **Reported clusters of activation across omnibus meta-analysis.** Here we report the coordinates, z value, p value, and cluster size of the cluster associated with the decisions phase while exploring or exploiting. Coordinates are reported in Montreal Neurological Institute (MNI) space.
